# Neutralizing vaccination against viral interleukin-6 from a primate rhadinovirus

**DOI:** 10.64898/2026.10.26.761420

**Authors:** Khoi N. Huynh, Joney LeSon, Sruthi Kalavacherla, Desmond P. Hartigan-O’Connor, Joseph Dutra, Ning Chin, Vivian Le, Denise R. Andal, Justin Fontaine, W. L. William Chang, Jesse D. Deere

## Abstract

Herpesviruses are ancient viruses that devote a significant fraction of their protein-coding potential to interference with host immunity. Diverse herpesviruses encode proteins with similarity to host MHC molecules, interferon regulatory factors, complement regulators, G-protein coupled receptors, chemokines, and cytokines. Certain gammaherpesviruses, including Kaposi sarcoma-associated herpesvirus (KSHV) and rhesus rhadinovirus (RRV), encode viral cytokines similar to host IL-6, called viral IL-6 (vIL-6). The potential of vIL-6 to act as a protective vaccine antigen against these rhadinoviruses remains under-explored. We isolated a new RRV strain, which encodes virion surface proteins similar to those of previous isolates and a vIL-6 that is nearly identical. We found that, although unmodified vIL-6 is a poor immunogen eliciting low neutralizing titers, vIL-6-Fc fusion protein elicits high-titer responses after either one or two doses. We detected no adverse effects related to cytokine activity, nor interference with concomitantly administered vaccine against the RRV glycoprotein B. Our results demonstrate an approach for eliciting neutralizing antibodies against a highly conserved viral protein that contributes to rhadinovirus persistence and oncogenesis.

## 1 Introduction

As is true for other members of the *Herpesviridae*, Kaposi sarcoma-associated herpesvirus (KSHV), or human herpesvirus 8 (HHV-8), is likely most frequently transmitted via oral-fluid contact with mucosal surfaces. Emerging data suggest that the earliest host-virus interactions at these mucosal surfaces are essential to establishing primary infection, and that disrupting them can result in viral clearance. Subsequent to infection and systemic spread, KSHV utilizes a broad repertoire of strategies to circumvent host immunity and maintain persistent infection. One important strategy is to modulate immune responses using a viral homolog of the host IL-6 cytokine, called viral IL-6 (vIL-6).

Host IL-6 is a pleiotropic cytokine with broad pro- and anti-inflammatory effects regulating immune responses to insult. The cytokine is a key pro-inflammatory player in the acute-phase response but is also known as a “myokine” that helps to resolve inflammation and maintain tissue health. Chronically elevated IL-6 synthesis is most commonly associated with inflammation and autoimmunity (1). An anti-IL-6 receptor-neutralizing antibody, tocilizumab, was developed and has been used to treat several rheumatologic and autoimmune conditions, as well as multicentric Castleman disease (MCD) (2). The viral cytokine, vIL-6, contributes to KSHV-associated disease pathogenesis. Although this gene shares only limited homology with human IL-6, it activates the IL-6 signaling pathway by binding to the host gp130 receptor (without the need for binding to IL-6Rα) (3–5). vIL-6 is highly expressed in lytic replication and remains expressed at physiologically meaningful levels in latency (6, 7). vIL-6 stimulates expression of numerous pathogenic host factors such as vascular endothelial growth factor (VEGF) (8). vIL-6 also supports sustained KSHV transcription in monocytes and encourages differentiation of the cells into longer-lived macrophages (9). vIL-6 promotes oncogenesis via AKT pathway activation and STAT3 activation (10–13). STAT3 activation in turn drives angiogenesis via caveolin-1 downregulation and associated increases in VEGF expression (8, 14). Finally, vIL-6 contributes to KSHV persistence by inducing development of dysfunctional macrophages that are impaired in their ability to stimulate T-cells (9), and also impacts proper B cell development by increasing class-switching (15).

In previous work targeting the cytomegalovirus (CMV)-encoded homolog of IL-10 (cmvIL-10) through vaccination, we found that rhesus macaques that generated strong neutralizing antibodies to the cmvIL-10 protein were protected from horizontal challenge (16). This protection declined as anti-cmvIL-10 responses started to wane. Those results suggested that targeting viral homologs of host cytokines through vaccination has the potential to disrupt herpesvirus transmission or establishment of persistent infection.

Candidate vaccines against herpesviruses most commonly target virion-surface glycoproteins (17). These structural proteins are often required for viral entry into host cells through engagement of cellular receptors. KSHV has several key glycoproteins, including glycoprotein B (gB). As for other *Herpesviridae*, KSHV infection persists despite robust adaptive immune responses to these glycoproteins following infection. In addition, as for CMV, KSHV superinfection of previously exposed individuals is possible, showing that natural adaptive immunity to a very broad array of viral antigens does not prevent secondary infection (18). These findings suggest that targeting herpesvirus glycoproteins alone might not be sufficient to prevent infection and pathogenesis.

Our goal in this study was to use a rhesus rhadinovirus (RRV) nonhuman primate model of KSHV to determine the feasibility of eliciting immunity against both vIL-6 and gB. RRV is closely related to KSHV and similarly encodes a homolog of host IL-6 (19). Furthermore, it has been shown that following co-infection with SIV, some RRV isolates induce a hyperplastic B-cell multicentric lymphoproliferative disorder similar to that observed in AIDS patients infected with KSHV (20). In this study, we isolated and characterized a novel RRV strain from a cohort of seropositive rhesus macaques. We then assessed the immunogenicity of vaccines targeting gB and vIL-6.

## 2 Materials and Methods

### Animal use

This study was approved in advance by the University of California, Davis (UC Davis) Institutional Animal Care and Use Committee (IACUC) and was performed at the National Biomedical Research Institute (NBRI). Housing, diet, medical care, and all procedures were performed in accordance with UC Davis IACUC-established policies. UC Davis has an Animal Welfare Assurance on file with the NIH Office of Laboratory Animal Welfare and is fully accredited by the Association for the Assessment and Accreditation of Laboratory Animal Care International. The animals assigned to these studies were adult rhesus macaques (*Macaca mulatta*) of Indian origin.

### Virus isolation

Anticoagulated blood samples were collected from eight adult donor macaques in the NBRI conventional colony, which were RhCMV-seronegative and RRV-seropositive. PBMC were isolated using Lymphocyte Separation Medium (LSM). 5 × 10⁶ cells per donor were then incubated with 3 × 10^5^ primary rhesus dermal fibroblasts (PrFB) in one mL of primary fibroblast medium (DMEM + 20% FBS + 1X Pen/Strep + 10 mM HEPES). The cultures were expanded to T75 flasks over the course of 14 days before evaluation for CPE. Cells from flasks demonstrating apparent CPE were then scraped into the media, mixed and frozen at −80°C. DNA was extracted from 0.5 mL of this suspension using Qiagen DNeasy Blood and Tissue columns and then screened by conventional PCR for gB and vIL-6. Amplified bands were sequenced to confirm presence of gB and vIL-6 sequences. The crude stock from one macaque that was confirmed positive for CPE, gB DNA, and vIL-6 DNA was then further expanded in PrFb to generate a lab stock (RRVJ1.2).

### Genome sequencing

To generate material for genome sequencing, a lysate collected after four passages was clarified, concentrated over a 20% Sorbitol cushion, Benzonase treated to remove cellular DNA, and collected on another 20% Sorbitol cushion. Viral DNA was then isolated using a Zymo Mini quick DNA/RNA Plus column. vDNA was eluted with 30 µL of 1X IDTE and stored at 4°C. Isolated viral DNA was submitted for Illumina short-read sequencing. Sequencing reads were assembled *de novo* using SPAdes genome assembler v4.2.0 (21), and the largest contig was found using NCBI BLASTn to have high identity to RRV isolate 17577. Shorter contigs averaged 260bp in length and were excluded from downstream analysis.

Open reading frames (ORFs) were predicted from the assembled contig with Prodigal v2.6.3 and GeneMarkS v4.28 (22, 23). Predicted ORFs from each tool were filtered to those encoding proteins with length ≥100 amino acids. The resulting predictions were compared and consolidated to identify candidate ORFs, which were queried using NCBI BLASTp, restricted to *Herpesviridae*, to identify potential homologs. To identify additional candidate ORFs in regions with sequence similarity to RRV isolate 17577 that were not represented among the initial filtered ORFs, predicted ORFs encoding proteins of 50-99 amino acids were subsequently included and evaluated.

To improve resolution of repetitive regions of the genome, 3.2 μg of viral DNA were submitted for long-read sequencing on the PacBio Revio platform using standard PacBio SMRT Cell configurations at the UC Davis DNA Technologies Core. HiFi PacBio reads were mapped to RRV isolate 17577 using Minimap2 v2.30 (24). Reads containing unique sequences flanking internal and terminal repeat regions were extracted. Repeat sequences were examined directly from the extracted reads.

### High multiplicity-of-infection (MOI) growth curve

PrFB were seeded in 96-well plates at 10^4^ cells/well and infected the following day with RRVJ1.2 at an MOI of 0.5. Virus was allowed to adsorb for 1 hour at 37°C with gentle rocking every 15 minutes. Following adsorption the viral inoculum was removed, cells were washed with HBSS, and fresh growth medium was added. Cell-associated and extracellular virus fractions were collected separately at each time point over a 5-day time course. For collection of the cell-associated fraction, cells were treated with trypsin and resuspended in the same volume of complete growth medium as had been present in the well. Supernatants were clarified with centrifugation at 600 × g for 5 minutes. Samples were then stored at −80°C until infectious virus was titered by plaque assay.

### Multi-step growth curve

PrFB were seeded in 24-well plates at 4 × 10^5^ cells/well and infected the following day with RRVJ1.2 at a MOI of 0.05. Virus was allowed to adsorb for 1 hour at 37°C with gentle rocking every 15 minutes. Following adsorption the viral inoculum was removed, cells were washed with HBSS, and fresh growth medium was added. Cell-associated and extracellular virus fractions were collected separately at each time point over an 11-day time course. For collection of the cell-associated fraction, cells were treated with trypsin and resuspended in complete growth medium. Supernatants were clarified with centrifugation at 600 × g for 5 minutes. Samples were then stored at −80°C until infectious virus was titered by plaque assay.

### Plaque assay

Infectious RRV titers from growth-curve samples were determined using plaque assays on PrFB. PrFB were seeded in 6-well plates at 5 × 10^5^ cells per well one day before infection. Samples were serially diluted in growth medium, from 10^-1^ to 10^-4^ and 1 mL of the appropriate dilution was added to each well. A mock-infected well receiving growth medium without virus was included as a negative control.

Virus was allowed to adsorb for 1 hour at 37°C with gentle rocking every 15 minutes. Following adsorption, viral inoculum was removed, cells were washed with HBSS and 2 mL of overlay medium were added to each well. The final overlay contained 0.75% methylcellulose in 1x DMEM supplemented with 1x penicillin/streptomycin, 5% FBS, and sodium bicarbonate. Plates were allowed to rest for 30 minutes at room temperature to allow cellulose to set. Plates were then returned to 37°C with 5% CO2 and were monitored for plaque formation.

At day 11 post infection, cells were fixed with 4% paraformaldehyde (PFA) in PBS. Following fixation, plates were aspirated, and cell monolayers were stained with 0.1% Crystal Violet solution. Plates were rinsed, allowed to dry and plaques were counted. Viral titers were calculated and reported as plaque-forming units per milliliter (PFU/mL).

*Viral replication kinetics* at both low and high MOIs were modeled using an Erlang-distributed multiple-eclipse-phase framework, implemented in R (deSolve). In this model, susceptible target cells (T) become infected via contact with free virus (V), progressing through three successive eclipse stages (*E*_1_, *E*_2_, *E*_3_) at a transition rate of *k* = *n*/*τ_E_*, where *τ_E_* is the mean total eclipse duration. Cells exiting the final eclipse stage become productively infected cells (I) that produce cell-associated virus (C) at a rate *p*. These cells die at a rate *δ*, while cell-associated virus is released into the supernatant at a rate *q*. Clearance of cell-associated and free virus occurs at fixed rates *c_C_* and *c_V_*, respectively. The incidence of infection is given by *β*(*V*/vol)*T* + *κIT*, where *β* is the infection rate constant and *κ* represents direct cell-to-cell spread. The model incorporates an adsorption loss term accounting for the consumption of free virus during initial infection. Model parameters were jointly estimated by minimizing the penalized negative log-likelihood across all conditions, explicitly handling left- and interval-censored titer measurements using a robust bounded multistart optimization approach (L-BFGS-B). Confidence in the eclipse phase duration was assessed by fixing *τ_E_* across a defined grid and re-optimizing all remaining nuisance parameters to generate a likelihood profile.

### Phylograms and cladograms

Whole-genome pairwise distances were calculated using Mash v2.3 (25) with a k-mer size of 21 and a sketch size of 10,000. The resulting distance matrix was used to construct a Neighbor-Joining (NJ) tree in R version 4.5.0 (26), which was then rooted on the betaherpesvirus outgroups. For the conserved core-gene analyses, protein sequences across six essential herpesvirus loci were aligned using MUSCLE (27). Maximum likelihood (ML) phylogenetic reconstructions and bootstrap consensus trees were subsequently generated using MEGA12 (28).

### Vaccine preparation

gB and vIL-6 mRNAs were produced by *in-vitro* transcription using N1-methylpseudouridine and capped using CleanCap, then packaged into GenVoy nanoparticles on the Precision Nanosystems Ignite instrument. mRNA/LNP vaccines were validated before use by transduction of Expi293 cells and testing for synthesized protein by western blot or ELISA. 25 mcg of each mRNA were administered intramuscularly per dose. Recombinant vIL-6-Fc protein was generated by transfection of Expi293 cells. Supernatants were collected after centrifugation (3200 × g for 20 min at 4 °C) and filtered through a 0.45-micron PES filter. Protein A resin was pre-packed in a borosilicate glass column and equilibrated with 15 column volumes (CV) of wash buffer. The supernatant was loaded over the column three times and washed with 10 CV of wash buffer. vIL-6- Fc was eluted with 5 CV 100 mM citrate, pH 3.5, into tubes containing 1 M Tris, pH 8. Elution fractions containing protein were checked using SDS-PAGE with and without DTT as the reducing agent in a Mini-PROTEAN TGX gel, quantified by Bradford assay (Bio-Rad Laboratories), and frozen at –20°C. vIL-6-Fc was mixed with an equal volume of AddaSO3 adjuvant (InvivoGen) before intramuscular administration.

### gB ELISA

gB-specific antibodies were measured by indirect ELISA. Plates were coated overnight at 4°C with gB-expressing cell lysate at 0.5 mcg/mL, using 100 µL per well. Plates were washed with PBST and blocked with Thermo Scientific SuperBlock. Serum samples were collected at multiple time points from rhesus macaques vaccinated with either gB or gB+vIL-6 mRNA vaccines. These samples were then serially diluted and added to the gB-coated wells. Bound antibodies were detected using goat anti-human IgG Fc-HRP at 1:5,000. Plates were developed with TMB, stopped with 2N H_2_SO_4_ and absorbance was measured at 450nm.

### vIL-6-Fc ELISA

vIL-6-Fc specific antibodies were measured by indirect ELISA. Plates were coated overnight at 4°C with purified recombinant vIL-6-Fc at 1 mcg/mL, using 100 µL per well. Plates were washed with PBST and blocked with Thermo Scientific SuperBlock. Serum samples were collected at multiple time points. These samples were then serially diluted and added to the vIL-6- Fc-coated wells. Bound antibodies were detected using goat anti-human IgG Fab-HRP at 1:5,000. Plates were developed with TMB, stopped with 2N H_2_SO_4_ and absorbance was measured at 450 nm.

### vIL-6-Fc neutralization

Serum samples collected from vIL-6 mRNA vaccinated rhesus macaques were heat-inactivated at 56°C for 30 minutes and diluted in HEK-Blue test medium. Diluted serum was incubated with recombinant vIL-6-Fc for 5 hours at 37°C prior to addition of the mixture to HEK-Blue IL-6 reporter cells (InvivoGen). Each well received 100 µL of serum+vIL-6-Fc mixture and 5 × 10^4^ reporter cells in 100 µL of test medium. Cells were incubated overnight at 37°C with 5% CO_2_. Controls included cells incubated with test medium plus serum alone (no vIL-6) and test medium supplemented with vIL-6-Fc. All conditions were assayed in duplicate. Following incubation, plates were centrifuged at 600 × g for 5 minutes, and 20 µL of supernatant were transferred to 180 µL of QUANTI-Blue reagent (Invivogen). Plates were incubated for 2 hours at 37°C and absorbance was measured at 620 nm to quantify reporter cell activity.

### Viral neutralization assay

Serum samples collected from gB vaccinated macaques were heat- inactivated at 56°C for 30 minutes, diluted in primary fibroblast medium, and mixed with RRV stock. The samples were incubated at 37°C for 1 hour to allow for neutralization of the virus, and then the mixture was added to PrFB seeded in 24-well plates at 0.8 × 10^5^ cells/well in 1 ml of primary fibroblast medium. Virus was allowed to adsorb for 2 hours at 37°C with gentle rocking every 15 minutes. Following adsorption, the viral inoculum was removed, cells were washed with 1x PBS, 1 ml of growth media was added, and the cultures were returned to 37°C with 5% CO_2_. Four days after inoculation, the cultures were assessed for cytopathic effect (CPE).

## 3 Results

### Characterization of a rhesus rhadinovirus isolate, RRVJ1.2

PBMCs were isolated from eight macaques at the NBRI that were seronegative for rhesus cytomegalovirus (RhCMV) and seropositive for RRV. The cells were co-cultured with PrFBs for two weeks, at which point two cultures with apparent cytopathic effect (CPE) were harvested and designated viral passage 1. The identity of the virus was confirmed by amplification and sequencing of the glycoprotein B and vIL-6 genes. One isolate, designated RRVJ1.2, was further expanded on fibroblasts before collection on a sorbitol cushion and genome sequencing.

Inoculation of passage-4 virus onto new PrFB monolayers produced progressive cytopathic effects (CPE) that became increasingly apparent during 11 days of observation (Figure 1A, left). Control cultures maintained monolayers of intact, spindle-shaped cells. Virus-exposed monolayers were instead progressively disrupted, with loss of normal morphology, rounding and aggregation, and increasing numbers of refractile or detached cells and cell-free areas. Dense cellular aggregates were observed after about 8 or 9 days. Some aggregates appeared to be multinucleated syncytia, but multinucleation could not be definitively established. Such changes were consistently more pronounced in inoculated cultures than controls, supporting progressive CPE rather than nonspecific deterioration.

**FIGURE 1.**
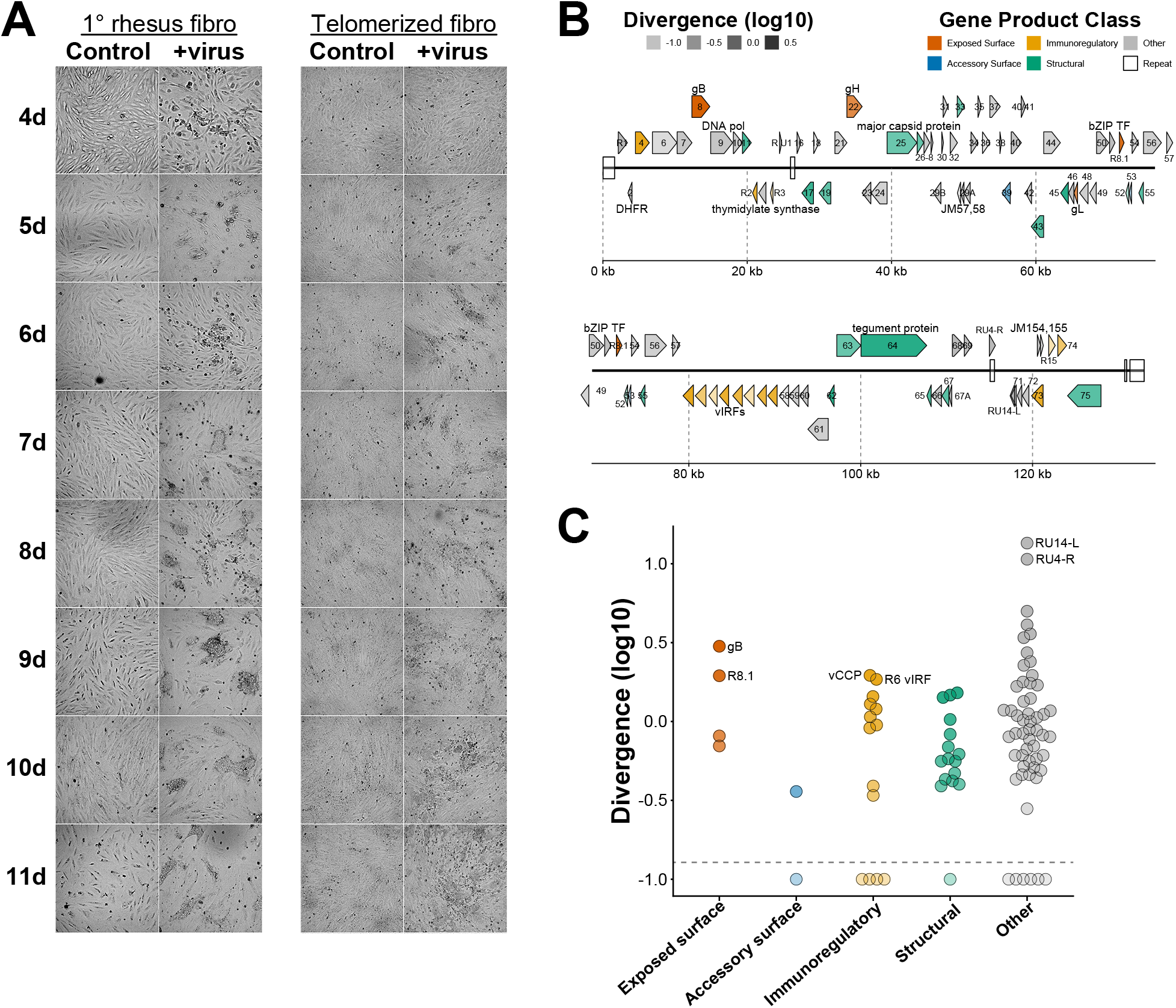
Basic characterization of RRVJ1.2. **(A)** Progression of cytopathic effect in PrFB (left) or telomerized rhesus fibroblasts (right) over days 4–11 post-infection. Images were captured at 10× magnification. **(B)** Map of annotated ORFs and repeat regions in the genome. ORFs are color-coded based on the functional class of their gene products, with arrows oriented in direction of transcription. Shading intensity of each ORF is in proportion to amino acid sequence divergence from RRV isolate 17577, where divergence is calculated as 100.1% – % sequence identity. **(C)** Distribution of amino acid sequence divergence of RRVJ1.2 ORFs from RRV isolate 17577 grouped by class of gene product. Divergence is plotted on a base-ten logarithmic scale.

Inoculation of telomerized rhesus fibroblasts (telos) caused more slowly progressive CPE (Figure 1A, right). Control telo cultures maintained a stable, uniform monolayer throughout the experiment. In virus-inoculated telos, the first changes at days 4-5 were subtle, consisting of scattered refractile cells and dark punctate cells on an intact background. Areas of cell rounding, increased refractility, and granular aggregation then became increasingly prominent. By days 9 to 11, these foci expanded and coalesced into dark aggregates accompanied by clearings in the monolayer consistent with cytolysis.

The viral genome was colinear with and very similar in coding content to that of other RRV isolates (Figure 1B). The maximum divergence from RRV isolate 17577 (29) was seen for RRV-unique (RU) ORFs on both strands of the GC-rich internal repeat region at approximate positions 115,000 to 118,000 (>10% non-identity). While exploring homologies in this region, we observed with surprise that the rightward ORF, RU4-R, is highly self-similar when translated in any of three rightward reading frames (Supplementary Figure 1). Exposed virion surface proteins, including gB and R8.1, were also somewhat divergent (2.9% and 1.8% non-identical; Figure 1C). Most predicted proteins, including vIL-6, were at least 99% identical to homologous proteins from isolate 17577 (Supplementary Table 1).

### Growth kinetics and tropism of RRVJ1.2

To characterize the replication kinetics of RRVJ1.2, PrFBs were infected at an MOI of 0.5 and infectious virus associated with the cells or released into the culture supernatant was quantified by plaque assay for the next four days (Figure 2A). Infectious virus in the supernatant increased substantially by 24 hours after infection and reached approximately 7.3 × 10^3^ PFU/mL by day 4, whereas cell-associated virus initially declined and remained below ∼1.7 × 10^3^ PFU/mL. At all time points after day 0, infectious virus was recovered predominantly from the supernatant. Thus, after infection at high multiplicity, RRVJ1.2 undergoes productive replication in rhesus fibroblasts and releases the majority of infectious progeny into the extracellular medium, consistent with efficient production and egress of cell-free virus.

**FIGURE 2.**
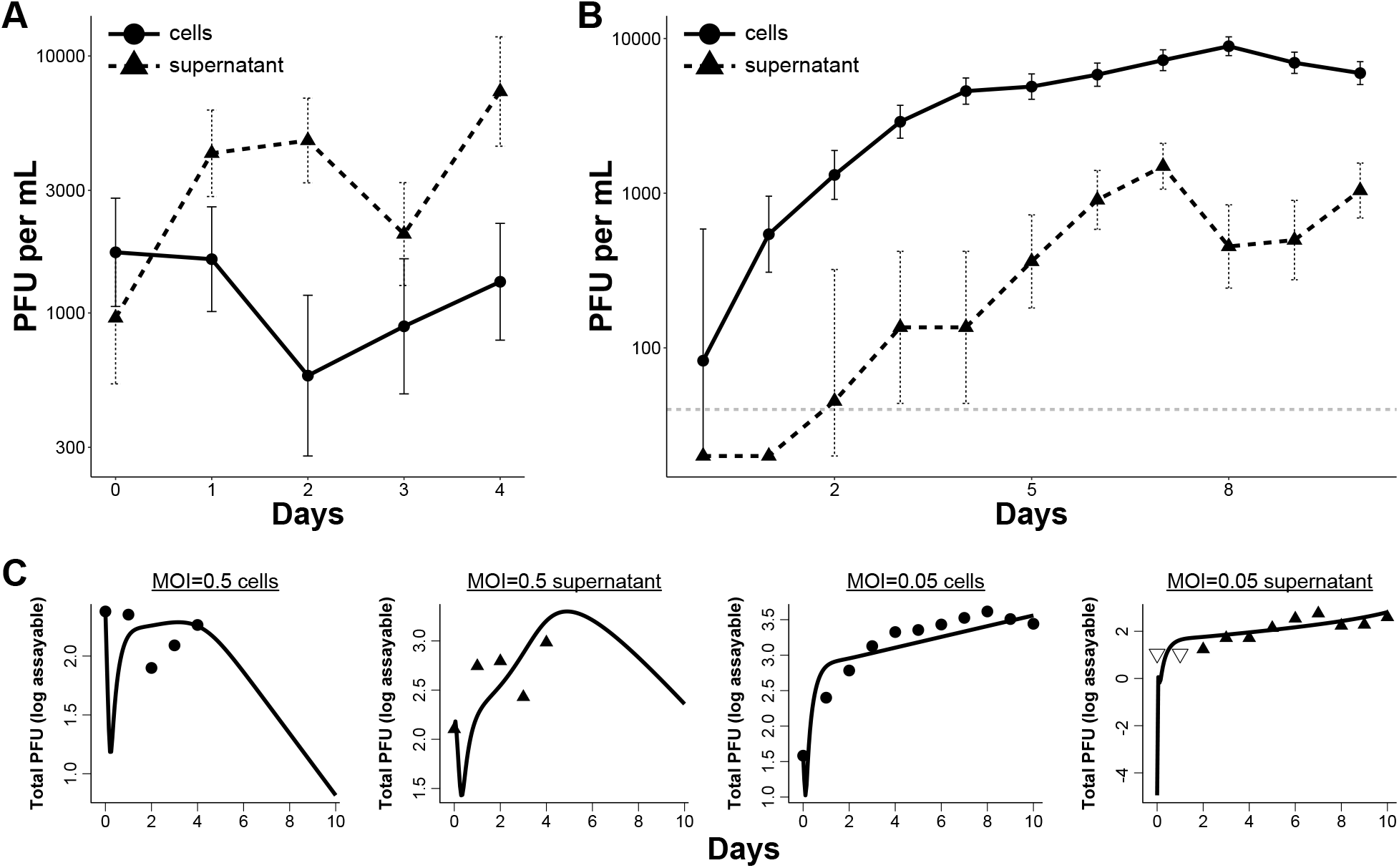
**(A)** Growth kinetics for RRVJ1.2 after inoculation of PrFBs at MOI = 0.5 for four days after inoculation. **(B)** Growth kinetics for RRVJ1.2 after inoculation of PrFBs at MOI = 0.05 for ten days after inoculation. Cell-associated and extracellular infectious virus titers were quantified by plaque assay. **(C)** Model-predicted viral kinetics (solid lines) from an Erlang-distributed eclipse phase model fitted to observed PFU production in cell and supernatant compartments (points).

To examine multistep growth kinetics, the fibroblasts were next infected at an MOI of 0.05 and infectious virus in the cell-associated and supernatant fractions quantified by plaque assay for ten days (Figure 2B). Cell-associated virus increased progressively from approximately 80 PFU/mL at day 0 to a peak of ∼9,000 PFU/mL at day 8, followed by modest declines over subsequent days.

Supernatant virus was below the limit of assay detection for the first two time points and was first quantifiable on day 2. Titers increased thereafter to a peak of approximately 1,500 PFU/mL one week after infection. At every measured time point, cell-associated infectious virus exceeded supernatant virus, by approximately one order of magnitude at day 7.

Thus, the distribution of infectious progeny between cells and supernatant differed between higher- and lower-multiplicity infections. At an MOI of 0.5, infectious RRVJ1.2 accumulated predominantly in supernatant, consistent with efficient release of virions from productively infected fibroblasts. In contrast, during multistep growth initiated at an MOI of 0.05, progeny virus was predominantly cell- associated. Possibly this difference reflects a greater contribution of cell-to-cell propagation after inoculation at low MOI, which is a mode of spread previously demonstrated for RRV and other herpesviruses (30–36), while infection at higher MOI reveals the capacity of synchronously infected cells to release virions extracellularly.

To quantitatively explore the replication dynamics, we fitted a mathematical (Erlang) model of viral kinetics jointly to the high- and low-MOI data (Figure 2C). The model tracked target cell infection, three eclipse phases, and the different trajectories of virus production in the cell-associated and supernatant fractions. Initial states were conditioned on the baseline titers (day 0) measured experimentally after washing the inoculum off of the target cells. The fitted model inferred a mean eclipse duration of 17.9 hours (0.75 days), followed by a productive cellular lifespan of 1.65 days, yielding a mean total time from infection to cell death of 2.4 days. The model suggested a transient cell-associated viral residence time of only 1.5 hours prior to extracellular release. Therefore, the growth-curve data suggest that RRVJ1.2 undergoes a protracted eclipse phase and exports mature virions efficiently, supporting the idea that accumulation of cell-associated virus observed during multi-step (low MOI) growth is driven by local cell-to-cell propagation rather than defective virion egress.

### Phylogenetic relationships

To evaluate the relationship of RRVJ1.2 to other gammaherpesviruses at the whole-genome level, pairwise genomic distances were calculated using Mash (k-mer size 21, sketch size 10,000) and used to construct a Neighbor-Joining (NJ) tree, which was subsequently rooted on the betaherpesvirus outgroups Human cytomegalovirus (HCMV) and Rhesus cytomegalovirus (RhCMV). In the resulting phylogram (Figure 3A), the catarrhine primate rhadinoviruses, including KSHV, RFHV, and the RRV strains, form a contiguous phylogenetic cluster as expected. However, unexpectedly, the murid (MHV-68) and New World primate (HVS) rhadinoviruses are segregated from the Old World rhadinoviruses, clustering instead with the macavirus, AlHV-1. Furthermore, the lymphocryptovirus, Epstein-Barr virus, branches as a sibling taxon to the KSHV/RFHV/RRV clade. The whole-genome k-mer profiles, which are sensitive to bulk genomic content and gene acquisition, thus produce an unexpected phylogeny.

**FIGURE 3.**
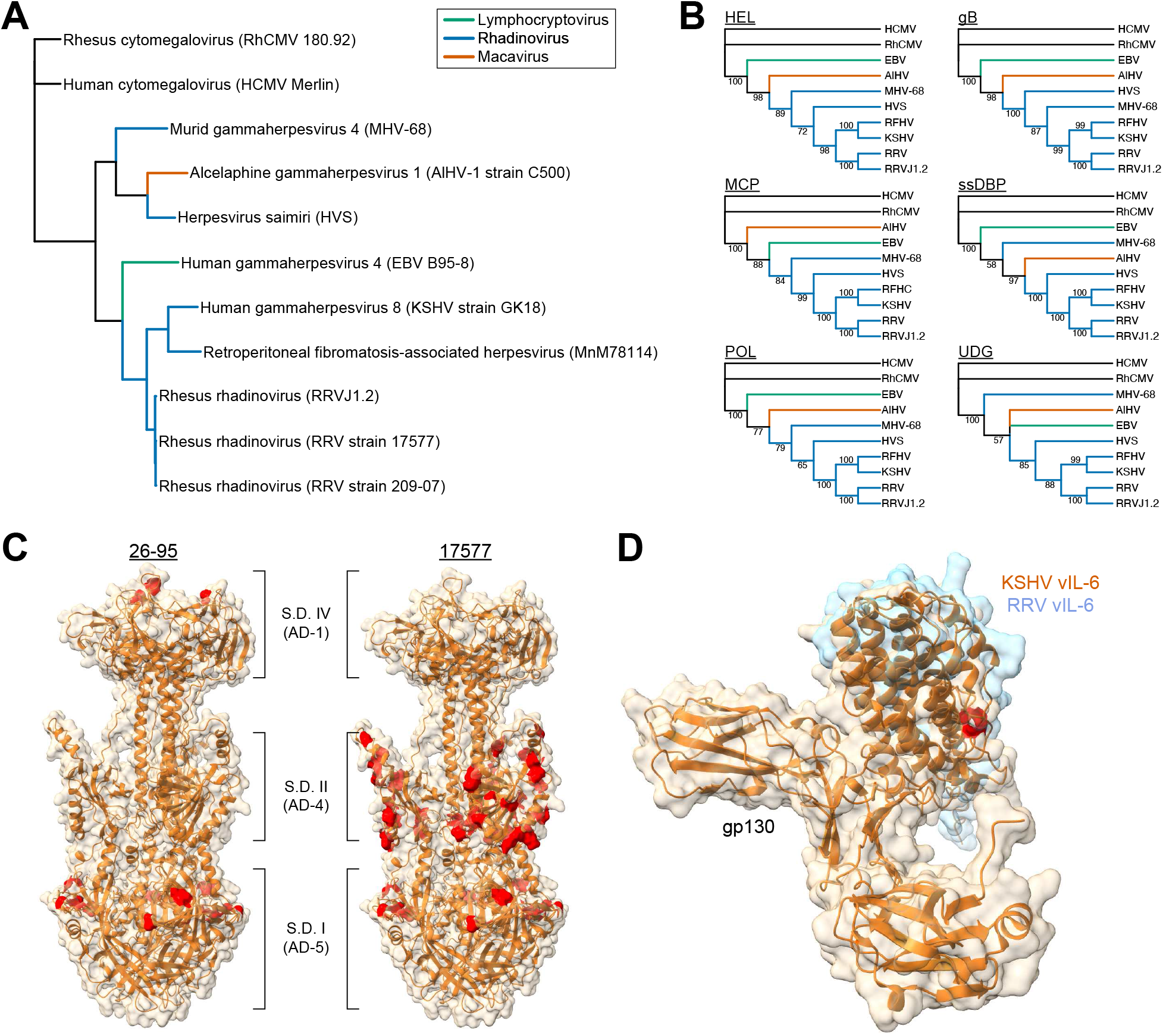
Phylogenetic and structural analysis of RRVJ1.2. **(A)** Neighbor-joining phylogram constructed from whole-genome pairwise k-mer distances produced by *mash* and rooted on betaherpesvirus outgroups. **(B)** Maximum-likelihood cladograms based on protein alignments of six conserved core genes (HEL, gB, MCP, ssDBP, POL, and UDG), from bootstrap consensus trees. **(C)** AlphaFold-predicted model of the RRVJ1.2 gB trimer. Residues differing from strains 26-95 (left) and 17577 (right) are highlighted, which demonstrates localization to established antigenic domains. **(D)** Predicted structure of the RRVJ1.2 vIL-6 protein matched using ChimeraX *matchmaker* to the KSHV vIL-6:gp130 complex, showing that variant amino acids are located distal to the vIL-6:gp130 interaction site.

Cladograms based on conserved core-gene alignments (Figure 3B) produced more conventional trees. Maximum likelihood (ML) phylogenetic reconstructions were performed on MUSCLE protein alignments across six essential herpesvirus loci: helicase (HEL), glycoprotein B (gB), major capsid protein (MCP), single-stranded DNA-binding protein (ssDBP), DNA polymerase (POL), and uracil-DNA glycosylase (UDG). Topologies were evaluated using bootstrap consensus trees. RRVJ1.2 clustered exclusively with RRV strain 17577 with 100% support. The core-gene topologies restored the expected monophyletic organization of the catarrhine rhadinoviruses. In two cases, however, MHV-68 failed to cluster with the primate rhadinoviruses (Figure 3B, trees for ssDBP and UDG).

This unexpected variation indicates that in the rodent lineage these genes either have experienced divergent evolutionary pressures or possess different ancestral histories compared to their primate counterparts.

Because gB and vIL-6 are both potential vaccine targets, we examined the differences between these proteins encoded by RRVJ1.2 vs. previously reported strains 26-95 and 17577 (Supplementary Figure 2). We first constructed a model of the RRVJ1.2 gB extracellular trimer domain (residues 58-692) using AlphaFold, which was reported with metrics pTM = 0.8 and ipTM = 0.8, suggesting that the predicted fold could be similar to the true structure. The model had a structure that was reminiscent of reported HCMV glycoprotein B structures such as 5C6T (37), with apparently identical subdomains arranged similarly. We then highlighted the residues in this model that differ from those in gB proteins from RRV strains 26-95 or 17577 (Figure 3C). This analysis showed that the differing residues were all located within antigenic domains (ADs) AD-1 and AD-5, for the contrast with strain 26-95 gB, or in AD-4 and AD-5, for the contrast with strain 17577. Localization of the amino-acid variation to these important targets of neutralizing antibodies suggests antigenic drift.

The predicted RRVJ1.2 vIL-6 amino-acid sequence has two variant amino acids compared to the same protein in strains 26-95 and 17577 (Supplementary Figure 2). We modeled the protein structure by matching the AlphaFold-predicted structure of the RRVJ1.2 vIL-6 protein to the experimentally determined, gp130-complexed structure of the KSHV vIL-6 protein, 1I1R (38). We found that the first variant (V31A) is located within a flexible segment without coordinates in the 1I1R structure, while the second (A164V) is located distal to the gp130 interaction site (Figure 3D). Therefore neither mutation is likely to impact the vIL-6:gp130 interaction.

### Immune responses to vIL-6 mRNA/LNPs or vIL-6-Fc fusion protein

The NBRI maintains a unique breeding colony of macaques (SPF level 2+) free of RRV. Macaques outside of this SPF level 2+ colony are uniformly seropositive by adolescence (Figure 4A). Interestingly, only a fraction of these macaques produce antibodies against vIL-6 (Figure 4B). There is a non-significant trend to lower anti-gB antibody titers in macaques reactive to vIL-6—suggesting the possibility that vIL-6 neutralization can assist in controlling viral replication (Figure 4C).

**FIGURE 4.**
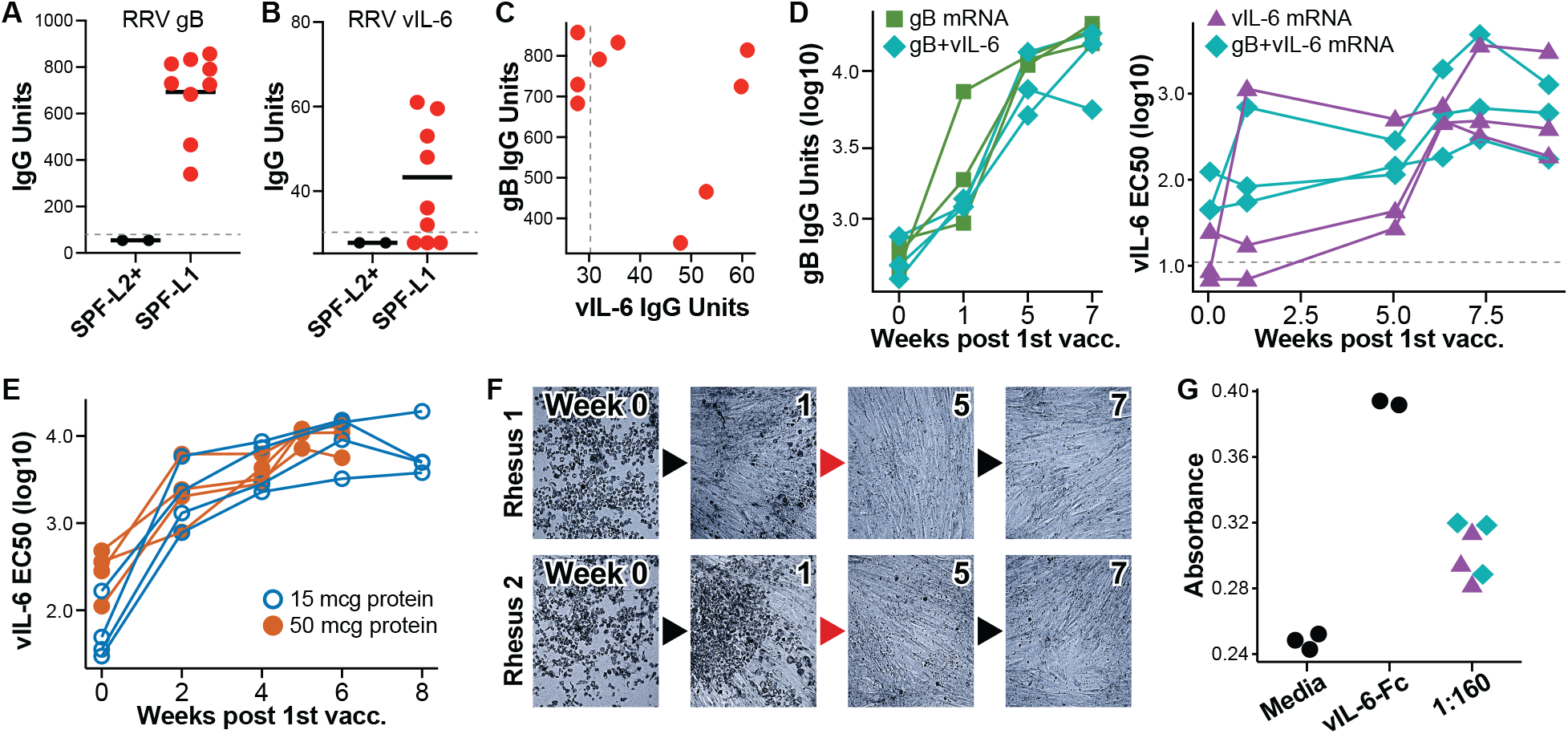
mRNA/LNP or novel protein immunogens boost immunity against vIL-6. **(A)** Anti-gB antibody responses among macaques from NBRI’s RRV-free SPF Level-2+ colony (left); or SPF Level-1 colony (right). **(B)** 6/9 RRV+ macaques have anti-vIL-6 titers. **(C)** Trend to lower anti-gB titers among macaques with antibody to vIL-6. **(D)** mRNA/LNP vaccines generate robust bAb responses to RRV gB and vIL-6, which are also neutralizing (F-G). **(E)** Vaccination with vIL-6-Fc protein using AddaSO3 adjuvant generates more intense bAb responses to vIL-6. **(F)** At left, RRVJ1.2 mixed with pre-vacc. sera lyses cells but mixed with post-boost sera (right) is non-infectious. **(G)** Neutralization of vIL-6 functional activity seen seven weeks after priming vaccination, assessed using HEK-Blue IL-6 Cells.

To test immunogenicity of mRNA/LNP vaccines expressing RRV gB or vIL-6, we administered the vaccines twice (0 and 5 weeks) to nine seropositive recipients (Figure 4D). Three macaques received mRNA expressing gB only, three mRNA expressing vIL-6 only, and three received both vaccines.

The results demonstrated satisfactory boosting of antibody titers in 6/6 gB vaccine recipients and 6/6 vIL-6 recipients (Figure 4D); however, the maximal EC50 titers attained against vIL-6 were ∼1:1,000, which are relatively low compared to those routinely seen for other mRNA vaccine targets. We hypothesized that poor protein secretion might cause the suboptimal response to mRNA-encoded vIL-6. We therefore next fused the vIL-6 coding sequence to a macaque IgG1 Fc region, which facilitates both protein export and purification. We produced and purified these proteins and then administered them with Adda-SO3 adjuvant at low or high doses (15 or 50 mcg per vaccination, respectively; Figure 4E). This alternative approach resulted in over ten-fold higher maximal EC50 titers, >1:10,000. Interestingly, vaccination with 15 mcg protein was equally effective to vaccination with 50 mcg protein. No adverse effects or clinical signs of vIL-6-Fc administration were detectable by cage-side monitoring or complete blood counts.

Interestingly, the anti-gB antibody present before vaccination (Figure 4A) was non neutralizing (Fig. 2F, left). However, the vaccine-induced antibodies were weakly neutralizing after the priming dose and strongly neutralizing by 1w after boost (Figure 4F). There was no interference between the two mRNA/LNP vaccines when administered together. Post-vaccination anti-vIL-6 antisera were also neutralizing when assayed at high dilution (Figure 4G), while at lower dilutions the assay appeared compromised by residual host IL-6 activity.

## 4 Discussion

Prototype rhesus rhadinovirus (RRV) isolates were previously isolated from captive macaque colonies, notably strains 17577 and 26-95 (19, 39). These isolates have been shown to exhibit broad tropism *in vivo*, infecting primary B lymphocytes, monocytes, endothelial cells, and epithelial tissues. They have the potential for pathogenesis, inducing a hyperplastic B-cell lymphoproliferative disorder resembling multicentric Castleman disease when introduced to SIV-infected macaques (20). Our isolate, RRVJ1.2, mirrors the previously demonstrated tropism while revealing subtle host-cell dependencies. RRVJ1.2 undergoes rapid lytic replication in primary rhesus fibroblasts, generating robust CPE and high viral yields, while infection of telomerized rhesus fibroblasts results in attenuated, slowly progressive CPE. The sequence of RRVJ1.2 is highly collinear with strains 17577 and 26-95. However, localized divergence in surface glycoproteins gB (2.9% non-identical) and R8.1 (1.8% non-identical) is evidence of antigenic drift among circulating macaque isolates.

We observed different partitioning of RRVJ1.2 progeny into cell-associated and supernatant fractions depending on the multiplicity of infection. At high MOI (0.5), infection proceeded almost synchronously and yielded rapid release of cell-free virus into the supernatant. Conversely, low-MOI infection (0.05) led to predominantly cell-associated viral titers that exceeded those in the supernatant by nearly an order of magnitude. Our kinetic model resolved a mean eclipse duration of 17.9 hours (0.75 days), a productive lifespan of 1.65 days, and a remarkably short cell-associated residence time of only 1.5 hours prior to egress. If this model is accurate then (because virion export is rapid) the accumulation of cell-associated virus during low-MOI infection likely reflects direct cell-to-cell spread across the monolayer (30–36).

These viral dynamics align with the results of a quantitative study that documented different cell-associated and cell-free viral dynamics following high-versus low-MOI HSV infection (40). The investigators found that high MOI produced a shorter eclipse phase, earlier viral egress, and faster infected-cell death. Low-MOI infection was dominated by cell-to-cell transmission: direct spread accounted for most new infections under low-MOI conditions, with cell-free infection making only a small contribution. The authors concluded that their results demonstrated different cell-associated and cell-free viral dynamics following infection at high versus low MOI. Indeed, frequent cell-to-cell transmission is typical across the *Herpesviridae*, which use tegument-mediated assembly (e.g., ORF52 in RRV) and routing to virological synapses to shield virions from extracellular clearance (30, 34, 35). By quantitatively assessing the eclipse phase and egress kinetics, our model confirms that RRV utilizes rapid intracellular processing coupled with efficient cell-to-cell dissemination to sustain infection.

Targeting herpesvirus-encoded cytokines is a promising vaccine strategy, which could add to the limited effectiveness seen so far (17). It seems apparent that vaccine strategies focused only on neutralizing antibodies will be insufficient for protection, perhaps due to a combination of cell-to-cell transmission and virus-encoded immunomodulators such as vIL-6. Proof-of-concept for cytokine-targeted vaccination was established in the rhesus cytomegalovirus model, where we showed that neutralizing antibodies against cmvIL-10 significantly reduced horizontal virus transmission and altered long-term immune persistence (16). However, no prior vaccine study has reported targeting of KSHV or RRV vIL-6.

We found that while mRNA/LNP vaccines expressing native vIL-6 produce modest antibody titers (∼1:1,000 EC₅₀), fusion of vIL-6 to an Fc moiety dramatically enhances immunogenicity, yielding neutralizing antibody titers exceeding 1:10,000. Joint immunization against vIL-6 and gB is therefore possible and will achieve two aims: blocking entry via gB neutralization and simultaneously inhibiting immunomodulation driven by secreted vIL-6 (41). Our future experiments will therefore test the protective efficacy of vIL-6-Fc vaccination in macaque models against primary mucosal infection, viral shedding, latent reservoir establishment, or lymphoproliferative pathogenesis.

## 5 Data Availability Statement

The datasets generated for this study are presented in the article/Supplementary Material. Further inquiries can be directed to the corresponding author.

## 6 Author Contributions

KNH: Conceptualization, Formal analysis, Investigation, Methodology, Project administration, Resources, Supervision, Visualization, Writing – original draft, Writing – review & editing. JL: Formal analysis, Investigation, Resources, Visualization, Writing – original draft, Writing – review & editing. SK: Data curation, Formal analysis, Investigation, Visualization, Writing – original draft, Writing – review & editing. DPH-O: Formal analysis, Investigation, Visualization, Writing – original draft, Writing – review & editing. JD: Resources, Supervision, Writing – review & editing. NC: Data curation, Investigation, Supervision, Writing – review & editing. VL: Investigation, Writing – review & editing. DRA: Investigation, Supervision, Writing – review & editing. JF: Resources, Writing – review & editing. WLWC.: Investigation, Project administration, Resources, Supervision, Writing – review & editing. JDD: Conceptualization, Funding acquisition, Investigation, Methodology, Project administration, Supervision, Visualization, Writing – original draft, Writing – review & editing.

## 7 Funding

This work was supported with funding from the James B. Pendleton Charitable Trust; National Institute of Allergy and Infectious Diseases, National Institutes of Health (NIH) (Award No. R21 AI188878); and Office of the Director, NIH (Award Nos. P51 OD011107 and S10 OD036449).

## 8 Acknowledgments

We thank the veterinarians and staff, especially Miles Christensen, at the National Biomedical Research Institute. We thank Dennis Hartigan-O’Connor for advice throughout the project.

## 9 Conflict of Interest

WLWC and JD hold equity in Tendel Therapies Inc. The remaining authors declare that the research was conducted in the absence of any commercial or financial relationships that could be construed as a potential conflict of interest.

## Supplementary Material

### 1 Supplementary Figures and Tables

**Supplementary Figure 1.**
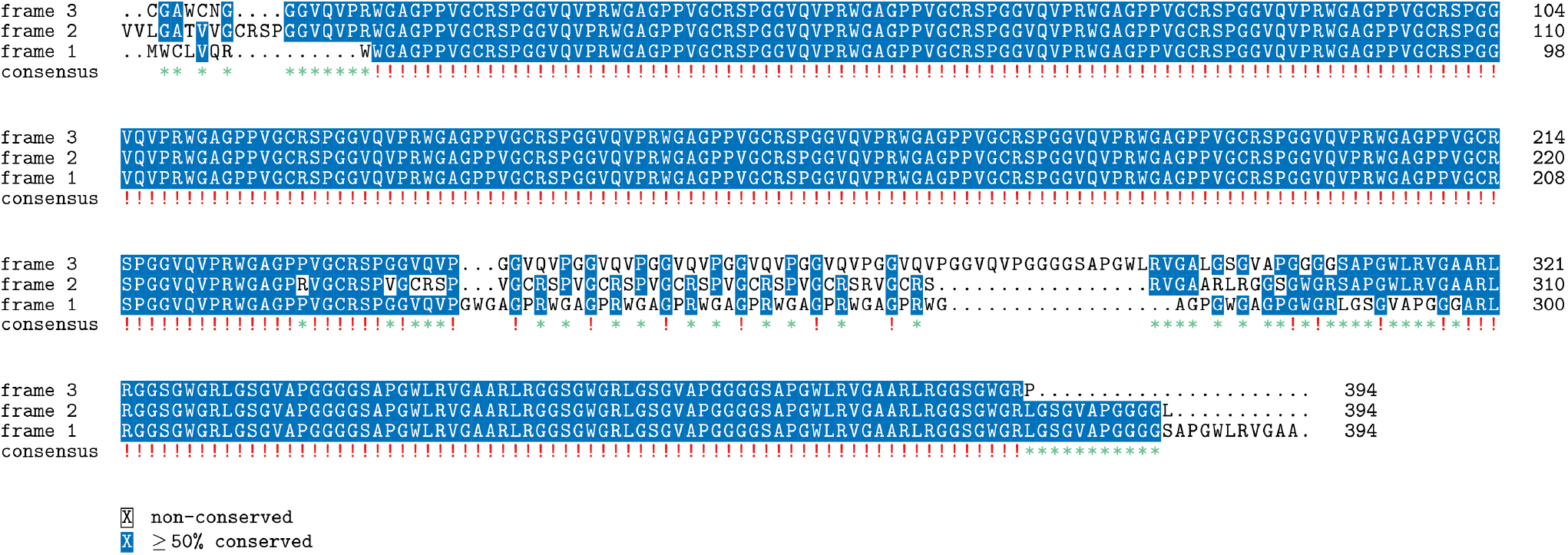
Extraordinary self-similarity of the RU4-R ORF when translated in any of three reading frames.

**Supplementary Figure 2.**
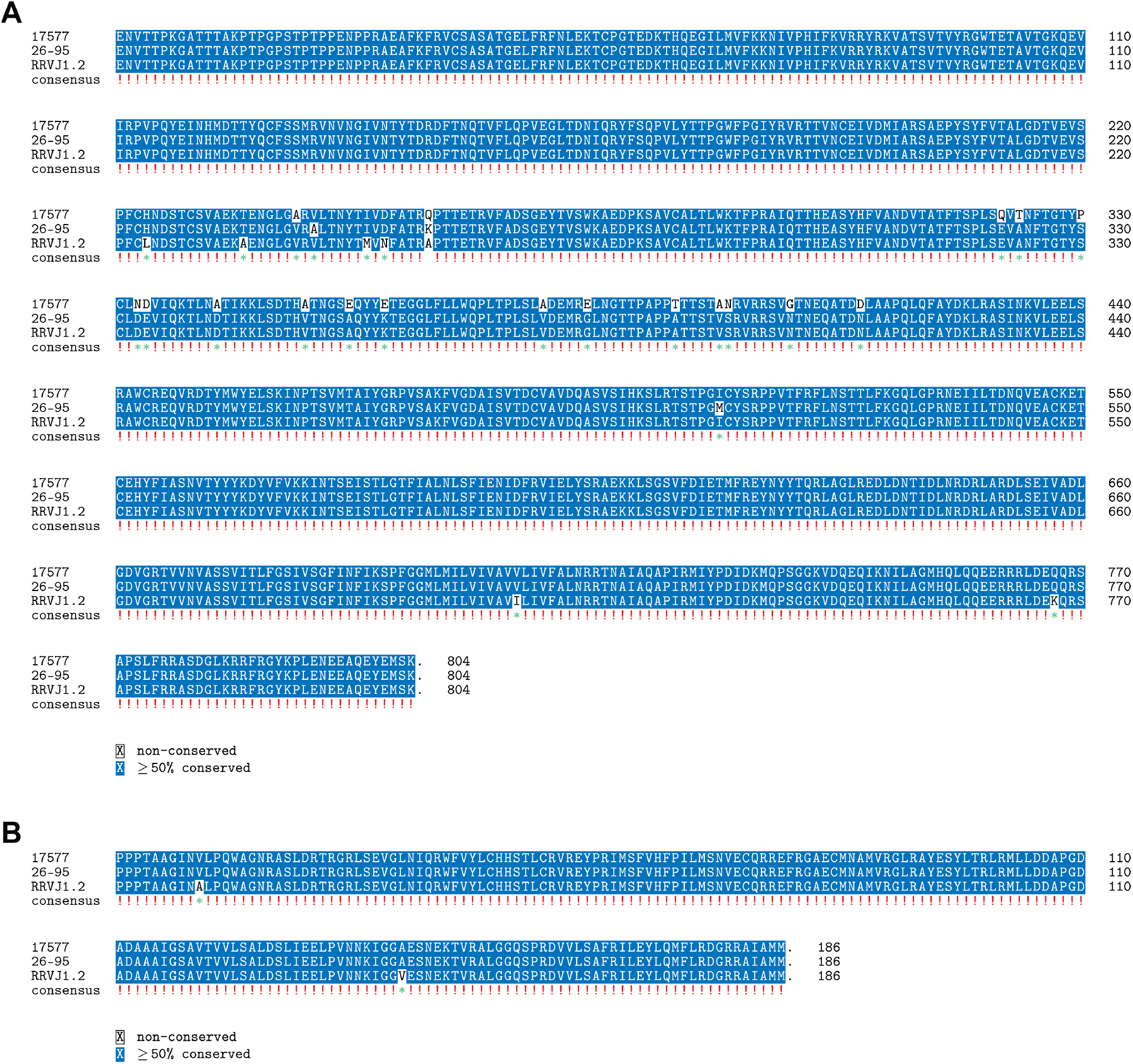
(A) Amino-acid alignment of the predicted glycoprotein B sequences from RRVJ1.2, 26-95, and 17577. (B) Amino-acid alignment of the predicted viral interleukin 6 sequences from RRVJ1.2, 26-95, and 17577.

**Supplementary Table 1.**
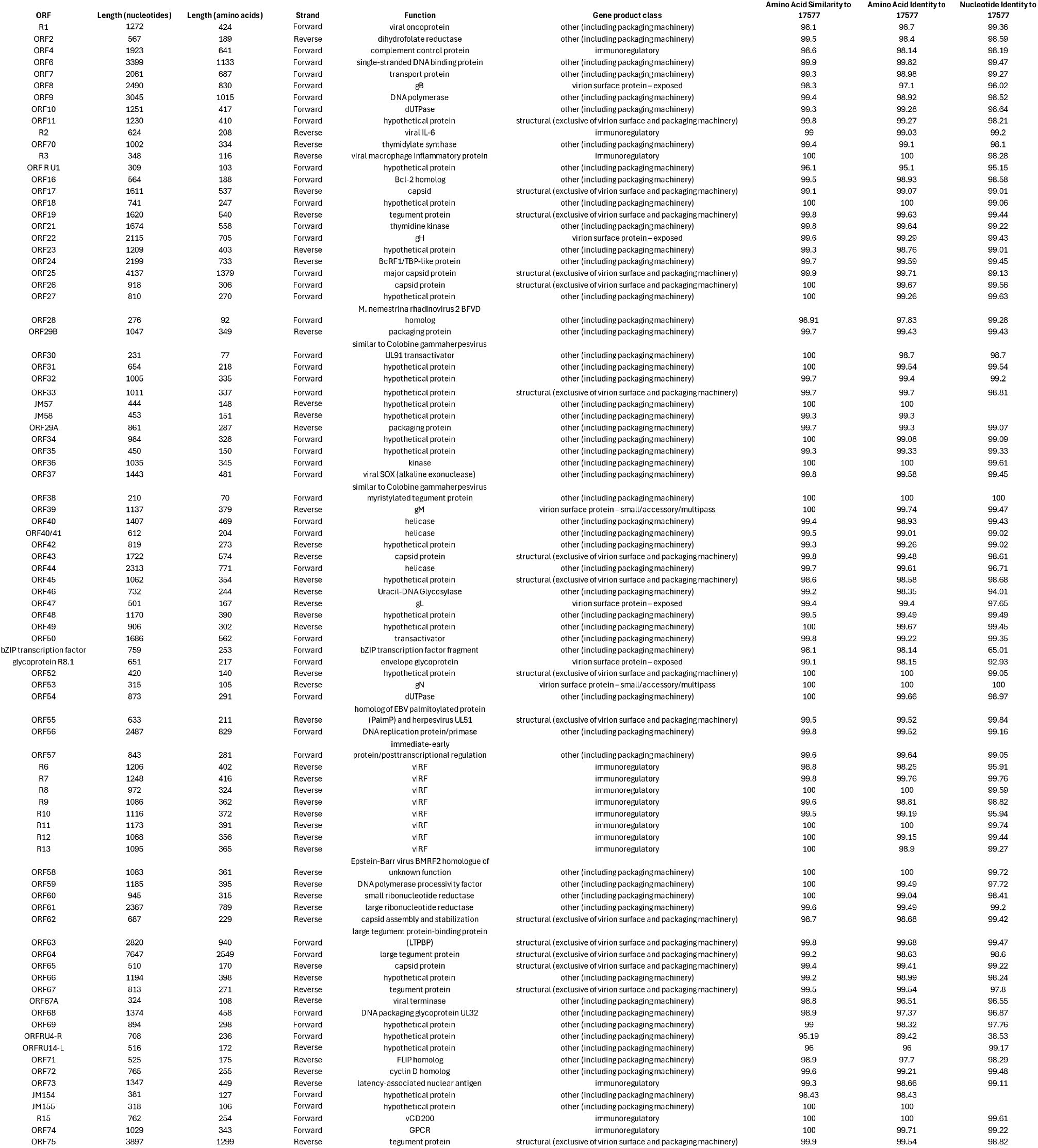
Prodigal- and GenemarkS-predicted open reading frames (ORFs) of RRVJ1.2 and their sequence identity and similarity to ORFs in RRV isolate 17577.

## References

1. Tanaka T, Narazaki M, Kishimoto T. IL-6 in Inflammation, Immunity, and Disease. Cold Spring Harbor Perspectives in Biology. 2014;6(10).

2. Rose-John S, Winthrop K, Calabrese L, Rose-John S, Winthrop K, Calabrese L. The role of IL-6 in host defence against infections: immunobiology and clinical implications. Nature Reviews Rheumatology. 2017;13(7).

3. Li H, Wang H, Nicholas J. Detection of direct binding of human herpesvirus 8-encoded interleukin-6 (vIL-6) to both gp130 and IL-6 receptor (IL-6R) and identification of amino acid residues of vIL-6 important for IL-6R-dependent and -independent signaling. J Virol. 2001;75(7):3325–34.

4. Molden J, Chang Y, You Y, Moore PS, Goldsmith MA. A Kaposi’s sarcoma-associated herpesvirus-encoded cytokine homolog (vIL-6) activates signaling through the shared gp130 receptor subunit. JBiolChem. 1997;272(31):19625–31.

5. Wan X, Wang H, Nicholas J. Human herpesvirus 8 interleukin-6 (vIL-6) signals through gp130 but has structural and receptor-binding properties distinct from those of human IL-6. J Virol. 1999;73(10):8268–78.

6. Chandriani S, Ganem D. Array-based transcript profiling and limiting-dilution reverse transcription-PCR analysis identify additional latent genes in Kaposi’s sarcoma-associated herpesvirus. J Virol. 2010;84(11):5565–73.

7. Sakakibara S, Tosato G. Viral interleukin-6: role in Kaposi’s sarcoma-associated herpesvirus: associated malignancies. J Interferon Cytokine Res. 2011;31(11):791–801.

8. Carbone A, Borok M, Damania B, Gloghini A, Polizzotto MN, Jayanthan RK, et al. Castleman disease. Nat Rev Dis Primers. 2021;7(1):84.

9. Shimoda M, Inagaki T, Davis RR, Merleev A, Tepper CG, Maverakis E, et al. Virally encoded interleukin-6 facilitates KSHV replication in monocytes and induction of dysfunctional macrophages. PLoS Pathog. 2023;19(10):e1011703.

10. Morris VA, Punjabi AS, Lagunoff M. Activation of Akt through gp130 receptor signaling is required for Kaposi’s sarcoma-associated herpesvirus-induced lymphatic reprogramming of endothelial cells. J Virol. 2008;82(17):8771–9.

11. Morris VA, Punjabi AS, Wells RC, Wittkopp CJ, Vart R, Lagunoff M. The KSHV viral IL-6 homolog is sufficient to induce blood to lymphatic endothelial cell differentiation. Virology. 2012;428(2):112–20.

12. Giffin L, West JA, Damania B. Kaposi’s Sarcoma-Associated Herpesvirus Interleukin-6 Modulates Endothelial Cell Movement by Upregulating Cellular Genes Involved in Migration. mBio. 2015;6(6):e01499–15.

13. Zhu X, Guo Y, Yao S, Yan Q, Xue M, Hao T, et al. Synergy between Kaposi’s sarcoma-associated herpesvirus (KSHV) vIL-6 and HIV-1 Nef protein in promotion of angiogenesis and oncogenesis: role of the AKT signaling pathway. Oncogene. 2014;33(15):1986–96.

14. Aoki Y, Jaffe ES, Chang Y, Jones K, Teruya-Feldstein J, Moore PS, et al. Angiogenesis and hematopoiesis induced by Kaposi’s sarcoma-associated herpesvirus-encoded interleukin-6. Blood. 1999;93(12):4034–43.

15. Rosario SA, Santiago GE, Mesri EA, Verdun RE. Kaposi’s Sarcoma-Associated Herpesvirus-Encoded Viral IL-6 (vIL-6) Enhances Immunoglobulin Class-Switch Recombination. Front Microbiol. 2018;9:3119.

16. Deere JD, William Chang WL, Villalobos A, Schmidt KA, Deshpande A, Castillo LD, et al. Neutralization of rhesus cytomegalovirus IL-10 reduces horizontal transmission and alters long-term immunity. Proceedings of the National Academy of Sciences of the United States of America. 2019;116(26):13036–41.

17. Moderna Announces Phase 3 Study of Investigational Cytomegalovirus (CMV) Vaccine Did Not Meet Primary Efficacy Endpoint [press release]. Moderna, October 22, 2025 2025.

18. Cornejo Castro EM, Marshall V, Lack J, Lurain K, Immonen T, Labo N, et al. Dual infection and recombination of Kaposi sarcoma herpesvirus revealed by whole-genome sequence analysis of effusion samples. Virus Evol. 2020;6(2):veaa047.

19. Desrosiers RC, Sasseville VG, Czajak SC, Zhang X, Mansfield KG, Kaur A, et al. A herpesvirus of rhesus monkeys related to the human Kaposi’s sarcoma-associated herpesvirus. J Virol. 1997;71(12):9764–9.

20. Wong SW, Bergquam EP, Swanson RM, Lee FW, Shiigi SM, Avery NA, et al. Induction of B cell hyperplasia in simian immunodeficiency virus-infected rhesus macaques with the simian homologue of Kaposi’s sarcoma-associated herpesvirus. J Exp Med. 1999;190(6):827– 40.

21. Bankevich A, Nurk S, Antipov D, Gurevich AA, Dvorkin M, Kulikov AS, et al. SPAdes: a new genome assembly algorithm and its applications to single-cell sequencing. J Comput Biol. 2012;19(5):455–77.

22. Besemer J, Lomsadze A, Borodovsky M. GeneMarkS: a self-training method for prediction of gene starts in microbial genomes. Implications for finding sequence motifs in regulatory regions. Nucleic Acids Res. 2001;29(12):2607–18.

23. Hyatt D, Chen GL, Locascio PF, Land ML, Larimer FW, Hauser LJ. Prodigal: prokaryotic gene recognition and translation initiation site identification. BMC Bioinformatics. 2010;11:119.

24. Li H. Minimap2: pairwise alignment for nucleotide sequences. Bioinformatics. 2018;34(18):3094–100.

25. Ondov BD, Treangen TJ, Melsted P, Mallonee AB, Bergman NH, Koren S, et al. Mash: fast genome and metagenome distance estimation using MinHash. Genome biology. 2016;17(1):132.

26. Team RDC. R: A Language and Environment for Statistical Computing. 2.13.0 ed. Vienna, Austria: R Foundation for Statistical Computing; 2011.

27. Edgar RC. Muscle5: High-accuracy alignment ensembles enable unbiased assessments of sequence homology and phylogeny. Nat Commun. 2022;13(1):6968.

28. Kumar S, Stecher G, Suleski M, Sanderford M, Sharma S, Tamura K. MEGA12: Molecular Evolutionary Genetic Analysis Version 12 for Adaptive and Green Computing. Mol Biol Evol. 2024;41(12).

29. Information NCfB. Accession No. NC_003401, Macacine herpesvirus 5, genome. In: (US) NLoM, editor. Bethesda (MD) 2018.

30. Anderson MS, Loftus MS, Kedes DH. Maturation and vesicle-mediated egress of primate gammaherpesvirus rhesus monkey rhadinovirus require inner tegument protein ORF52. J Virol. 2014;88(16):9111–28.

31. Black FL, Melnick JL. Microepidemiology of poliomyelitis and herpes-B infections: spread of the viruses within tissue cultures. J Immunol. 1955;74(3):236–42.

32. Campadelli-Fiume G, Roizman B. The egress of herpesviruses from cells: the unanswered questions. J Virol. 2006;80(13):6716–7; author replies 7–9.

33. Christian RT, Ludovici PP, Jeter WS. Cell-to-cell transmission of herpes simplex virus in primary human amnion cells. Proc Soc Exp Biol Med. 1971;138(3):1109–15.

34. Mothes W, Sherer NM, Jin J, Zhong P. Virus cell-to-cell transmission. J Virol. 2010;84(17):8360–8.

35. Sattentau Q. Avoiding the void: cell-to-cell spread of human viruses. Nat Rev Microbiol. 2008;6(11):815–26.

36. Tomescu C, Law WK, Kedes DH. Surface downregulation of major histocompatibility complex class I, PE-CAM, and ICAM-1 following de novo infection of endothelial cells with Kaposi’s sarcoma-associated herpesvirus. J Virol. 2003;77(17):9669–84.

37. Chandramouli S, Ciferri C, Nikitin PA, Calo S, Gerrein R, Balabanis K, et al. Structure of HCMV glycoprotein B in the postfusion conformation bound to a neutralizing human antibody. Nat Commun. 2015;6:8176.

38. Chow D, He X, Snow AL, Rose-John S, Garcia KC. Structure of an extracellular gp130 cytokine receptor signaling complex. Science. 2001;291(5511):2150–5.

39. Searles RP, Bergquam EP, Axthelm MK, Wong SW. Sequence and genomic analysis of a Rhesus macaque rhadinovirus with similarity to Kaposi’s sarcoma-associated herpesvirus/human herpesvirus 8. J Virol. 1999;73(4):3040–53.

40. Esmaeili S, Swan DA, Jerome KR, Schiffer JT, Walter M. Intracellular and extracellular dynamics of herpes simplex virus 1 DNA and infectious particles in epithelial and neuronal cells. bioRxiv. 2025.

41. Komaki S, Inagaki T, Kumar A, Izumiya Y. The Role of vIL-6 in KSHV-Mediated Immune Evasion and Tumorigenesis. Viruses. 2024;16(12).

